# Situation Models in the Brain are Used to Resolve Word References

**DOI:** 10.1101/2025.07.15.664757

**Authors:** Sarah Aliko, Eva Wittenberg, Jeremy I. Skipper, Steven L. Small

## Abstract

Resolving the meaning of a pronoun requires retrieving information about a unique individual (a referent) from many remembered experiences and inferences throughout sensory and cognitive modalities, from vision and audition to episodic memory and social cognition. Here, we hypothesize that the functional neuroanatomy of pronoun interpretation involves interaction across distributed neural networks, during which each referent activates its own unique sensorimotor neural fingerprint associated with these experiences.

To test this hypothesis, we collected fMRI data from 20 people watching a full-length movie, and developed a 3D branched convolutional neural network to distinguish movie characters from the fMRI signal across distributed sensorimotor regions. The same model distinguished the characters referenced by pronouns, using these same sensorimotor regions, supplemented by the hippocampus, precuneus, and medial prefrontal cortices.

This work has far-reaching implications for understanding the relations among the many domains and modalities of neural representation required for ecological language comprehension. In particular, the demonstration that situation models, implemented in distributed sensory-motor and association cortices, are involved in resolving reference, suggests a whole brain distribution for language processing.

## Introduction

In human cognition, few phenomena are restricted to a single system. For instance, while watching a movie, new information about each character leads to the formation of neural representations throughout the brain encoding how a character looks and sounds, how they act, and how they relate to other characters. These neural representations are assembled dynamically during the course of the movie, creating a unique multidimensional neural fingerprint for each character.

One process that is particular to human language is the use of open variables that are resolved by multiple sources of dynamic contextual information (Goldberg, 2006; Jackendoff, 2002). For instance, pronouns like *he* or *she* are placeholders that are filled by context (e.g., (Ariel, 2001; Brown-Schmidt et al., 2005). Take the following short snippet as an example: “Sarah wrote a paper about pronouns. She wanted to publish it.” In this, the placeholders are the pronouns “it” and “she”, filled in unambiguously by the context of the paper and Sarah. This process of pronoun resolution (or *anaphora resolution*), often more complex than in the example shown, provides a unique opportunity to study how the brain develops and then uses a novel and unique multi-domain representation for each person. We hypothesize that during both aspects of this process, brain activity associated with specific individuals is instantiated in a distributed neural network “fingerprint” that includes regions commonly associated with language and others not.

Although pronouns are pointers to meaning, they carry little meaning themselves (Nunberg, 1993). In English, gender, number, and case information unambiguously identify the referents of the two pronouns in the sentence “John went to pick up his daughter at school. He drove her home”. Through the gender information conveyed by the pronouns, we understand that ‘he’ refers to John and ‘her’ to his daughter. In general, when encountering pronouns in discourse, people are capable of quickly inferring to whom or what a pronoun refers (Greene et al., 1992). To accomplish this, linguistic and/or extralinguistic context are needed: For example, a listener will not know the referent (identity) of “she” in “she makes a mean negroni”, unless the sentence is contextualized.

Linguistic models have proposed that to identify a referent, a listener probes for relevant disambiguating information, including linguistic features such as gender, number, and case, syntactic and semantic biases, and discourse structure, as well as non-linguistic cues (Gordon et al., 1995; Wittenberg et al., 2021 for review). It has been proposed that such information forms a general contextual representation that is built up incrementally with each added word in a sentence (Altmann & Kamide, 2009), and that the human brain encodes this meaning representation to help the interpretation of a reference later (Wittenberg et al., 2021).

In cognitive science, these representations are sometimes called *situation models*, a term intended to capture the schematic nature of sensorimotor, emotional, procedural, spatial, and imagistic concepts of an event, character, location and action (Bartlett, 1932; Zwaan & Radvansky, 1998). Neuroscientists using brain imaging have found that this schematic knowledge is encoded in various brain areas, with early work suggesting narrative-specific activation in the dorsal medial portion of the prefrontal cortex (Yarkoni et al., 2008) and later more naturalistic experiments showing similar activity in what they describe as posterior medial cortex and medial prefrontal cortex (mPFC), as well as in the superior frontal gyrus (Baldassano et al., 2018).

Neuroimaging studies on narrative comprehension have further elaborated on the brain-behaviour relations in maintenance and retrieval of situation models. The role of the mPFC has been characterized as actively maintaining situation models for retrieval (Baldassano et al., 2018). The basis of this role may relate to the relationship of mPFC to other brain regions important for theory of mind (and “mentalising”) and for internal processing when individuals are not attending to sensory stimuli (i.e., the “default mode”), which are involved in decision-making and construction of imagery (Baetens et al., 2014; Euston et al., 2012; Isoda et al., 2013; Xu et al., 2005).

A recent review of the “default mode” and unifying model of its functional anatomy characterizes this set of brain regions as critical to coordinating neural processes for social cognition, self-reference, episodic memory, language, and semantic memory to create “a coherent internal narrative of our experiences” (Menon, 2023). The mechanisms of anaphora resolution require precisely the processes described in this model with neural representations throughout the brain requiring just this sort of coordination.

The generalised nature of situation models allows them to track event-based representations in sentences by linking antecedent concepts to newly encountered ones (Altmann & Kamide, 2009). Numerous linguistic studies have provided evidence for a role of situation models in processing of pronouns. Since pronouns on their own carry little information about the referent, their resolution relies on language features such as syntactic constraints, information structure, coherence relations, or verbal features (Kaiser et al., 2009; Kaiser & Trueswell, 2011a, 2011b) as well as discourse features and world knowledge, which together link the representations of the current and preceding context (McMillan et al., 2012). This overall process that can be characterized in part by applying the concept of a situation model.

Without focusing on pronouns *per se*, several imaging studies of comprehension have probed neural representations underlying situation models, and have proposed that their construction involves connecting sensorimotor, language and emotional representations. In the brain, this processing involves activation of brain regions commonly involved in episodic memory retrieval (Altmann & Ekves, 2019; Bergen et al., 2007; Zwaan, 2016; Zwaan et al., 1995; Zwaan & Radvansky, 1998; Zwaan et al., 2002). Other regions implicated in the retrieval of situation models during language processing support the theoretical position of Menon (2023) on the role of the “default mode”, and include the middle temporal gyrus (MTG), inferior frontal gyrus (IFG), and angular gyrus (AG) bilaterally (Baldassano et al., 2017; Whitney et al., 2009; Yarkoni et al., 2008). These regions also consistently appear in meta-analyses and task-based studies, thus suggesting a domain-general role in language processing rather than a specific role in reactivating situation models to support pronoun resolution (Aliko et al., 2023; Battistella et al., 2020; Xu et al., 2016).

Given the role of these regions in subsequent memory, particularly for informative sentences (Hasson et al., 2007), it is possible that these regions participate in referent-specific pronoun resolution through the coordinating role of the nodes that interact during the “default mode”. Although there is behavioural and linguistic evidence for the involvement of situation models in resolving pronoun referents, concrete evidence for this remains to be demonstrated in the brain.

Related neuroimaging research has suggested that activation of regions associated with content retrieval (e.g., memory of a person) might lead to activation of regions related to associated sensory or motor aspects of the remembered object (e.g., activation of the fusiform face area) (Nyberg et al., 2000; Woroch et al., 2019). Some such studies demonstrated that the activity patterns of visual representations of objects, places and faces are distributed across different regions in the visual cortex, depending on their category (Cichy et al., 2012). With respect to faces *per se*, other studies have been able to reconstruct specific human faces from their unique brain activity in visual regions (“fingerprints”) during free recall (Haxby et al., 2001; Norman et al., 2006; VanRullen & Reddy, 2019). These studies suggest (i) that the brain activates distinct perceptual patterns to process different visual objects, and (ii) that these patterns are reliably reactivated during retrieval. Although studies on context retrieval provide evidence for the distinct formation of representations and their retrieval pathways during processing of visual information, there remains a lack of evidence linking linguistic retrieval (e.g., pronouns) to imagistic simulations of situation models in the brain.

The current study tested three hypotheses about how the human brain uses situation models to encode information about discourse characters and then to access this information to perform anaphoric reference. These hypotheses are as follows:

(1) antecedent visual representations of a character activate sensory and motor regions to build unique situation models;
(2) these regions are later reactivated during pronominal reference in the absence of a character’s visual representation, thus linking the antecedent to the referent; and
(3) individual character references elicit mostly overlapping activity patterns, with few distinct voxel distributions that allow the brain to distinguish between character-specific situation models in memory. Whereas these overlapping representations are postulated to be in areas characteristically involved in memory encoding, model construction, and theory of mind (e.g., mPFC), the character-specific differences are predicted to be in and around sensorimotor regions, key areas for implementation of situation models.

To test these hypotheses, we used fMRI scans of 20 participants watching a full-length movie, and labelled faces and pronominal references of the two main characters. Taking advantage of recent methodological research on convolutional neural networks that have been able to explain complex task-dependent brain activity patterns in neuroimaging data (Carrito & Semin, 2019; Caucheteux et al., 2022; Devlin et al., 2019), we applied this method to anaphora resolution in the movie data. To accomplish this, we created a branched 3D convolutional neural network model trained on the fMRI BOLD signals evoked by the relevant visual and pronoun references. The model was first implemented to distinguish the character references in the visual and pronoun domains separately, and to then find shared activations of the visual and pronoun representations of each character. Finally, we performed guided backpropagation to identify the clusters of voxels that the model used to learn to distinguish each character reference.

## Methods

### Participants

We used fMRI data from 20 subjects (10 females, range of age 19-53 years, M = 27.7 years, SD = 10.1 years) from the Naturalistic Neuroimaging Dataset (NNDb) (Aliko et al., 2020). All participants watched the entire duration of “500 Days of Summer” during acquisition of functional MRI scans. None of the participants had previously seen this movie. “500 Days of Summer” is a 2009 romantic comedy-drama during which the main character Tom Hansen narrates his relationship with the second main character Summer Finn. Additional characters include friends and family members of Tom and Summer.

### Neuroimaging data

All participants watched the entire movie during BOLD acquisition on a 1.5T Siemens MAGNETOM Avanto machine. The following parameters were used: TR=1s, TE= 54.8 ms, flip angle = 75°, 40 interleaved slices, resolution = 3.2 mm isotropic. Participants took as few breaks as possible to allow for a more natural viewing. Anatomical T1-weighted scans were collected over 10 min (TR = 2.73 s, TE = 3.57 ms, 176 sagittal slices, resolution = 1.0 mm). Preprocessing was performed using the AFNI software’s afni_proc.py script, with the pre-processed data made available on OpenNeuro (https://openneuro.org/); for a more detailed account of data acquisition, quality, and preprocessing, please refer to the original NNDb article (Aliko et al., 2020).

### Face detection in movies

To detect character faces, we first selected the five main characters by filtering the five highest grossing actors in the movie from ‘The Movie Database’ website (themoviedb.org). We collected each actor’s images from Google Images, ensuring that the face of each actor was the main focus of the photo and that multiple angles of their face were included (e.g., left, right, up, down). Moreover, where possible, we downloaded images of actors from the specific movie, because movie makeup and costumes may significantly change the appearance of an actor. On average, 27.8 (SD=11.0) images were used for training the model for each actor.

We used an existing face recognition algorithm to detect each actor’s faces in the movie (github.com/ageitgey/face_recognition). The algorithm encodes and saves images of faces in 128 dimensions (i.e., lowering their dimensionality). To accelerate the face detection process, we used a multi-threaded batch approach for the encoding step and a multi-process approach for the detection step at each movie frame. We then divided the movie frame-by-frame using a python package called openCV (github.com/opencv/opencv). Frame counts were then transformed to seconds and results saved to a “.json” dictionary file of this format:

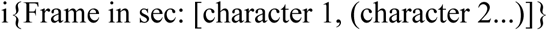

Predictions from the face recognition algorithm have associated error, which cannot be calculated unless all character faces are manually labelled frame-by-frame and compared to predictions. To reduce this error without the need for manual labelling, we only considered a character to be on screen if that character was predicted to be present in over 50% of the frames in one entire brain BOLD acquisition volume (1 sec). For instance, if within one functional volume there were 6/10 frames with predicted faces of Tom, then Tom’s face would be considered detected within that volume. Secondary characters were ignored here, even if they may have been present with one of the two main characters on screen. We avoided selecting any times where the main characters Tom and Summer appeared together on screen.

### Pronominal reference annotation

Movie audio signals were annotated using the Amazon AWS speech-to-text translator (aws.amazon.com/transcribe/). From the word transcripts, we selected the word timings for “500 Days of Summer” and identified all possible pronominal references to female and male referents (i.e., ”he/she”, ”his/hers”, ”him/her”). We labelled the pronoun onset as the start time of the subtitle containing the pronoun and used the functional volume containing the subtitle onset as the volume of interest. (Subtitles typically constituted short sentences spanning ∼1-2 acquisition volumes.) Finally, we manually labelled the character to which each pronoun referred and selected only pronominal instances referring to the two main leads in the movie (i.e., Tom and Summer). Although “she” and “he” pronouns were also used to refer to secondary characters, these instances were not of sufficient frequency for training a neural network. If a subtitle contained more than one pronoun, and these referred to different characters, these instances were also removed from the training data. In the end, the training data contained only cases where either Summer or Tom alone were being referenced through pronouns.

We next selected visual exemplars of each character that temporally overlapped with their pronoun references. We did this in two steps:

(1) First, we matched each pronoun occurrence with a visual image of Tom or Summer if the visual instance of the character occurred between 30 seconds and 2 minutes before the pronoun reference. The rationale for this is that we expected the situation model retrieved during the pronoun reference would be best represented by the closest antecedent character representation, i.e., that the visual reference of a character would be instances when a situation model was created (or updated) using perceptual information.
(2) Second, we ensured that instances of Tom and Summer were at least 5 seconds apart from each other. By doing so, we reduced potential overlaps between the two main characters either in the visual or auditory (pronoun) domains, particularly in scenes where they acted together, and the camera may have been switching between the two characters. All other pronoun samples were dropped, and unmatched visual samples were also removed. Although here we chose to focus on Tom and Summer, other characters were also occasionally on screen or speaking in a scene along with one of the two main characters.

### Data preprocessing and feature selection

To account for properties of the hemodynamic response function (HRF), we marked the onset of visual and pronoun representations 3 seconds after their occurrence, expecting the HRF to peak shortly thereafter (the typical lag is 5-7 sec from onset stimulus) (Yesilyurt et al., 2008). Next, we selected the functional (BOLD) volumes from 3 to 7 seconds after stimulus onset and averaged these to produce a less noisy signal centred around the theoretical peak of the HRF.

We identified more samples for Summer than those for Tom, and thus adjusted this imbalance by applying random oversampling with replacement on samples for Tom. This resulted in 21 samples per character for each participant (i.e., total of 420 samples for each character), with the final sample comprising 840 total samples of functional brain volumes (64×76×64 voxels) for each of the visual and pronoun samples (i.e., 1,680 brain volumes for the entire model).

For each participant, we masked the brain images by removing the voxels outside the brain and in the white matter and ventricles, and then computed a group mask to ensure all brain images had the same number of input features, which was a requirement for the input to a layer of a convolutional neural network (CNN) (Hashemi, 2019). Then we centred the samples to approximately have mean = 0 and standard deviation = 1, using the formula:

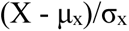

This preprocessing step helps the model learn and converge faster (Huang et al., 2020). To avoid overfitting, we randomly shuffled the functional volumes and labels. Labels for Tom and Summer were then converted to one-hot encoding (i.e., vectorised categorical labels) for input into the deep neural network.

### Model selection and training

The 3D branched convolutional neural network (CNN) takes as input the functional brain volumes of the matched visual and pronoun references of each character. Our CNN architecture was derived from an existing one proposed by Vu et al. (2020) for classifying four sensorimotor tasks in an fMRI experiment. Our model design included one visual branch and one pronoun branch to separately interrogate brain images representing visual and pronoun reference information. These two branches were then merged to identify any shared voxels of visual and pronoun referents.

As per the specifications detailed by Vu and colleagues, each branch consisted of 3 convolutional layers: Layer 1 (Conv1) had kernel size 7×7×7 and 8 filters, with a stride of 1; Layer 2 (Conv2) had kernel size 5×5×5 with 16 filters and stride of 1; and Layer 3 (Conv3) had kernel size 3×3×3 with 32 filters and strife of 1 (Vu et al., 2020). The first two convolutional layers were followed by an average pooling layer with stride 2 to reduce feature dimensions. Then we applied a flattening layer to vectorise convolved features to 1D, and finally by a fully connected layer with 128 nodes. We then added a further dropout layer with 50% retention probability to reduce overfitting. The two branches of the visual and pronoun streams were then concatenated with a further 50% retention probability dropout layer, and finally sent to a fully connected output layer with 2 nodes (i.e., classes). Figure 1 illustrates the structure of the model.

**Figure 1.**
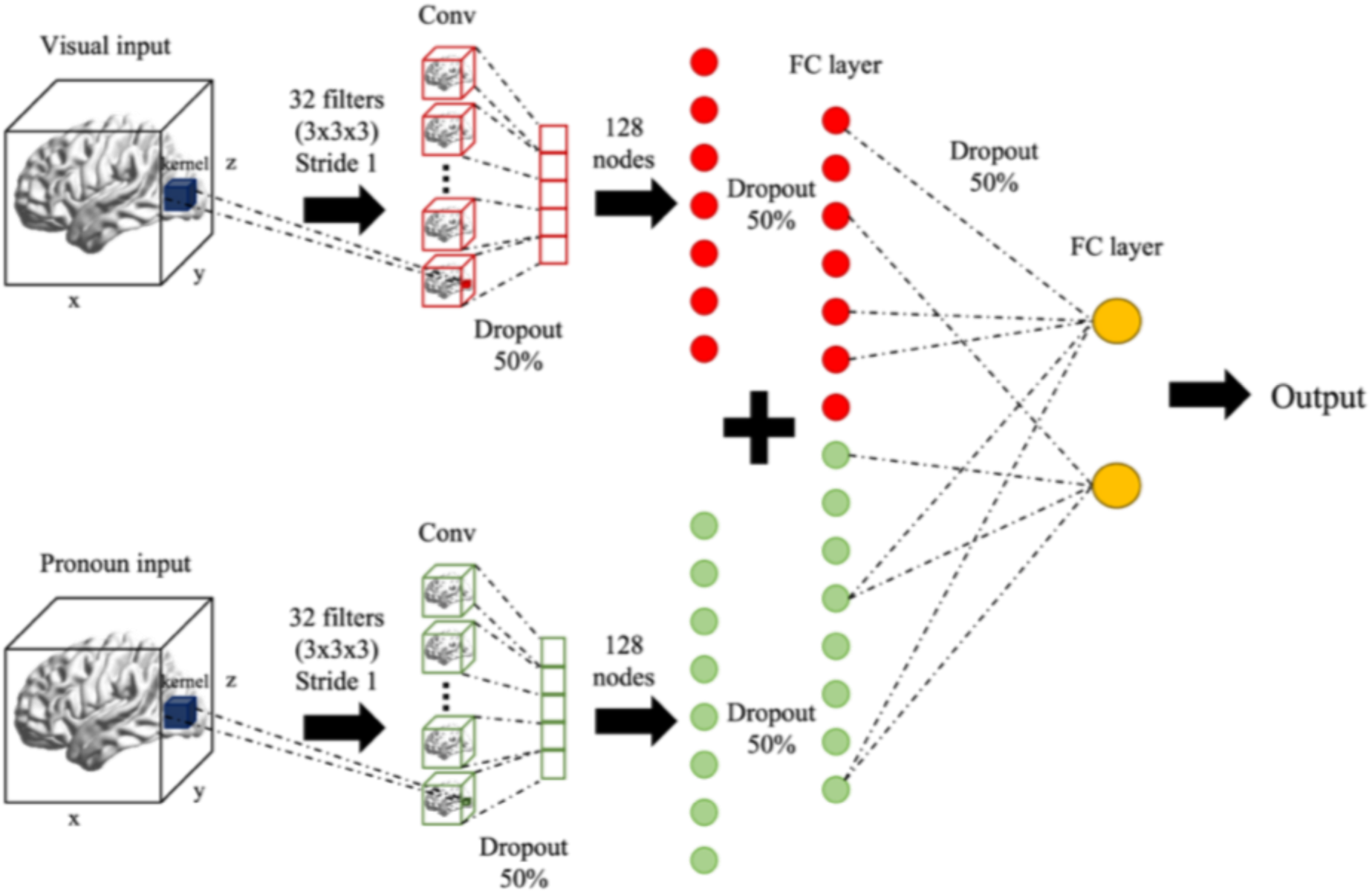
Final architecture and hyperparameters: In this final 3D CNN branched model, separate 3D volume inputs are fed into a visual and pronoun branch. The first layers of each branch consist of a 3×3×3 convolutional layer with 32 filters (stride 1), which are flattened and input into a 128 node fully connected layer. A 50% probability dropout reduces overfitting before merging the visual and pronoun branches. A final 50% dropout and fully connected layer (2 nodes) provide the output predictions.

Each convolutional layer and fully connected layers employed Rectified Linear Units (ReLU) activation functions, whereas the (fully connected) output layer used a ‘sigmoid’ activation function (Vu et al., 2020). Ridge regression (L2) regularisation was applied to the output layer activity to discourage overfitting.

The loss function for the model was binary cross-entropy, since we only had 2 classes (‘Tom’ and ‘Summer’). We applied a stochastic gradient descent (SGD) optimizer with initial learning rate (Lr_0_) of 10^-3^, which was step-decayed using the formula:

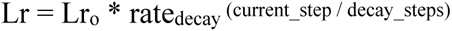

Where Lr is the new learning rate for a given step, rate of decay was 0.96 and decay steps were set to 100,000. Given that we had a small sample set that could be prone to overfitting in complex models, we tested 3 model complexities:

1. Three convolutional layers as in (Vu et al., 2020)
2. Two convolutional layers (removed Conv 1)
3. One convolutional layer (removed Conv 1 and Conv 2) and added 50% dropout after Conv 3 layer

Model performance was compared to two other convolutional neural networks, 2D and 3D CNN, using accuracy and loss as metrics. The 2D CNN comparison allows us to test whether using brain volumes instead of slices is better to identify relationships between voxel clusters, whereas the 3D CNN comparison enables examination of hyperparameters.

We did not compare our model with simpler models (e.g., support vector machines) because the design of two branching inputs in our model can only be accommodated by CNN models. The architectures and hyperparameters for the two comparison models are found in Supplementary Figures 1-3.

The data were split into 80% training and 20% testing. The performance of all three models were tested using 10-fold cross validation over 10 epochs on the training data and for each fold the accuracy and loss were also computed on the hold-out test data. The 3D bi-CNN was then trained on the entire data set (i.e., 1,680 pairs of samples) over 100 epochs with early stopping to reduce computational load.

### Saliency map visualisation

To visualise those voxels of the 3D images from which the model was learning, we computed saliency maps using vanilla guided backpropagation. Given an image *I* belonging to class *c* (either *Summer* or *Tom* here) the class score *S_c_(I)* can be approximated to a linear function using Taylor’s expansion rule, such that:

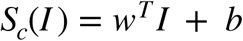

Where *w* is the weight of a voxel and *b* the bias term. Solving the derivative for *w*, we get the following:

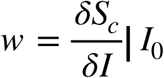

The collection of *w* for each voxel represents the saliency map of an image (Simonyan et al., 2013).

We selected the 3D volumes for *Summer* and *Tom* separately, ran each through vanilla guided backpropagation, and averaged the resulting maps for each image to obtain an overall map of each character representation. Due to the branched nature of our model, each character’s guided backpropagation resulted in a pronoun saliency map and a visual saliency map, which shared concatenation and output layers. We performed paired t-tests contrasting *Summer* vs *Tom* (visual + pronoun) and visual vs pronoun (*Summer* + *Tom*). We then thresholded each of the resulting saliency maps to the 95th percentile to maintain the strongest weights, thus preserving those voxels that contributed most to model learning. However, because we could not easily identify the specific sample index after the shuffling of the data, we could not separate the samples by participant (i.e., separate into 20 clusters), resulting in high degrees of freedom in the t-test.

## Results

### Model selection and performance

The 3D functional (BOLD signal) volumes were used as inputs into each of three convolutional neural networks (CNNs), all three containing a visual image branch and a pronoun branch (see Methods for details). The primary model was created by entering the 3D functional volumes to a CNN with decreasing kernel sizes over lower convolutional layers in each branch and then merging them into a fully connected layer, using the structure and hyperparameters of an existing 3D CNN architecture (Vu et al., 2020). The two comparison models were created as follows: (i) The 2D CNN was constructed by dividing the 3D volumes into 2D axial slices for input to a pretrained ResNet50 CNN for each branch and merging them into trainable fully connected layers. (ii) The 3D CNN was created by entering the 3D volumes into a CNN with increasing kernel sizes over lower convolutional layers in each branch and then merging them into a fully connected layer. With our primary model, we also tested three model complexities (as noted in Methods), using one, two, or three convolutional layers.

For the comparison models, (i) the transfer-learning model achieved an average validation accuracy of labelling Tom vs Summer both visually and through pronouns of 47.9% (SD = 10.7%), with M=0.72 (SD=0.05) loss over 10 folds. On the hold-out test data (20% of dataset), the model reached an average accuracy of 50.5% (SD=2.3%) and M=0.72 (SD=0.04) loss; and (ii) the 3D increasing kernel size CNN model reached an average validation accuracy of labelling Tom vs Summer both visually and through pronouns of 70.5% (SD = 5.3%), with M=0.64 (SD=0.01) loss over 10 folds. On the hold-out test data, the model reached an average accuracy of 70.1% (SD=3.4%) and M=0.64 (SD=0.01) loss.

Finally, we tested the primary model, the 3D decreasing kernel size CNN (Vu et al., 2020) at various complexities:

1. Three convolutional layers: reached an average validation accuracy of labelling Tom vs Summer both visually and through pronouns of 47.3% (SD = 10.9%) and an average validation loss of 0.69 (SD = 0.003). This model also achieved an average testing accuracy across folds of 56.8% (SD = 6.4%) and an average loss of 0.69 (SD = 0.0008).
2. Two convolutional layers: reached an average validation accuracy of labelling Tom vs Summer both visually and through pronouns of 71.7% (SD = 11.9%) and an average validation loss of 0.66 (SD = 0.01). This model also achieved an average testing accuracy across folds of 75.7% (SD = 6.1%) and an average loss of 0.66 (SD = 0.004).
3. One convolutional layer: reached an average validation accuracy of labelling Tom vs Summer both visually and through pronouns of 91.4% (SD = 2.0%) and an average validation loss of 0.34 (SD = 0.04). This model also achieved an average testing accuracy across folds of 92.6% (SD = 1.1%) and an average loss of 0.34 (SD = 0.02), indicating that it was stable across folds and appropriate for the task (i.e., not overfitting), unlike the former models. We named this final model ‘3D bi-CNN’. The training, validation, and testing accuracy per fold for these three final models are summarised in Figure 2.

**Figure 2.**
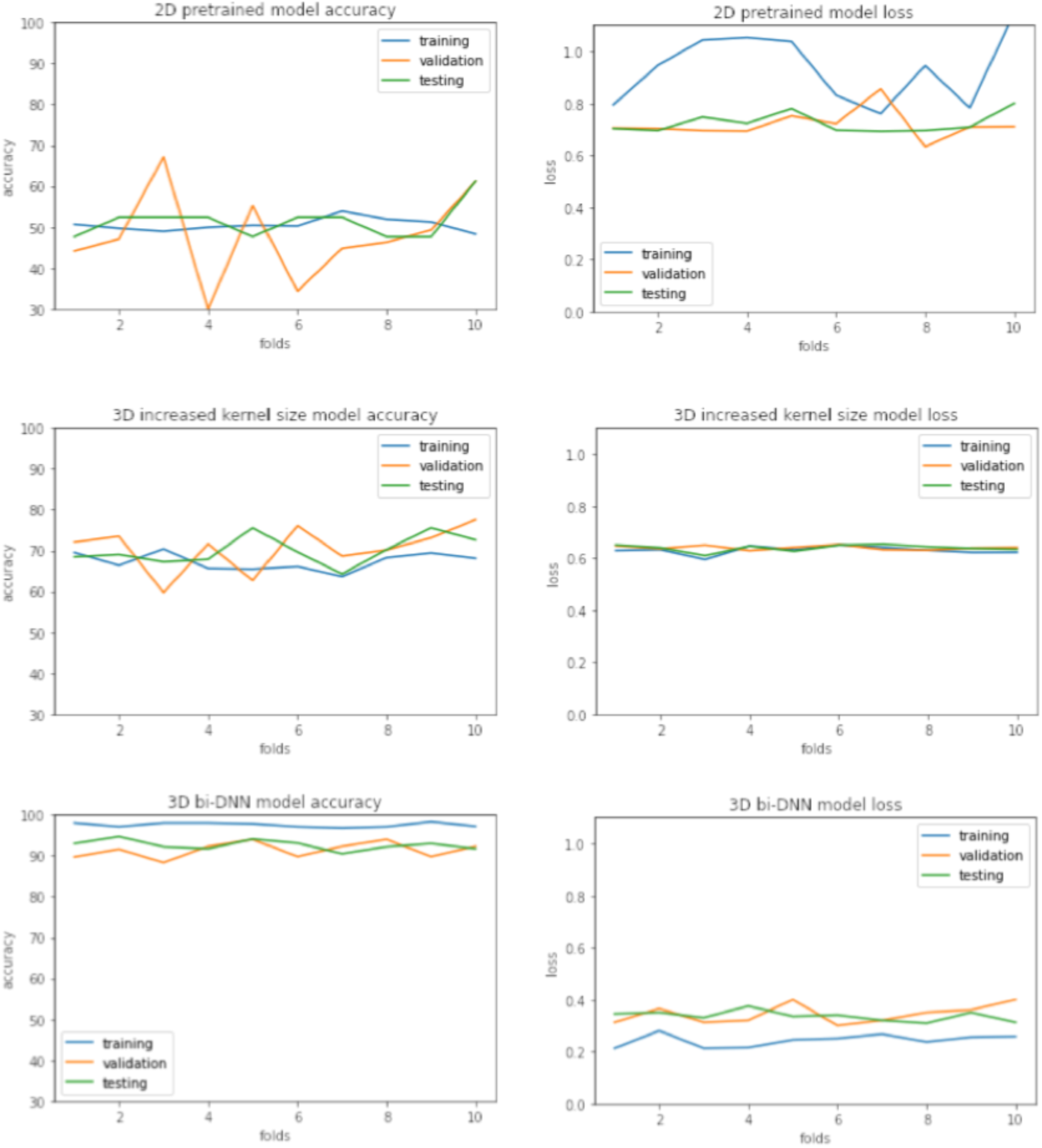
Plots of the accuracy (left column) and loss (right column) over 10 folds during cross validation. Over 10 folds: top row represents the RSN pre-trained model (47.9% accuracy, 0.72 loss); middle row is the 3D CNN with increasing kernel size (70.5% accuracy, 0.64 loss); bottom row is the final 1-CNN layer 3D CNN model (92.6% accuracy, 0.34 loss). The last model outperforms the others in terms of both increased accuracy and reduced error.

After selecting the 1-Conv-layer 3D bi-CNN as the best model, we trained it over 100 epochs on the entire dataset with early stopping to reduce the computational load.

### Saliency maps

We then focused on visualising the specific voxels the model used to distinguish pronoun references to Tom from those for Summer. To do this, we performed guided backpropagation as follows:

1) Run the image through the classifier (forward pass);
2) Calculate the gradient of the class with respect to the image as a derivative function;
3) Normalise gradients to values between 0 and 1.

This resulted in 420 maps for each of the two characters’ faces and 420 for each character’s pronoun referent. We performed paired t-tests contrasting (i) Summer vs Tom (visual + pronoun) and (ii) visual vs pronoun (Summer + Tom) over 840 samples each. We found all within-brain voxel comparisons to be significant (ps < 10^-5^), due to the large number of degrees of freedom.

For each of the four categories, the maps were averaged together into a single 3D volume and thresholded to 95^th^ percentile, to obtain only the most significant weights (or voxels). The results are shown in Figure 3. The most prominent regions used for statistical inference in the visual branch model were the primary visual cortex, higher-level visual regions (e.g., fusiform face area), superior temporal gyrus, parahippocampus, and temporal-parietal junction (TPJ) (Figure 3A). For the pronoun branch of the overall model, the main regions used for inference were primary visual cortex, higher-level visual areas, dmPFC, precuneus, and hippocampus (Figure 3B). The characters overlapped in over 38% of voxels from the visual model and over 57% in the pronoun model, and with differences primarily occurring in voxels around sensorimotor regions in both. Moreover, within each character’s maps, the visual and pronoun branches overlapped in 8% of voxels (for both Tom and Summer), primarily in the visual cortex and parahippocampal areas (Figure 3C). Visual references more commonly mapped to the superior temporal gyrus and occipital cortex, whereas pronoun references more commonly mapped to mPFC and precuneus.

**Figure 3:**
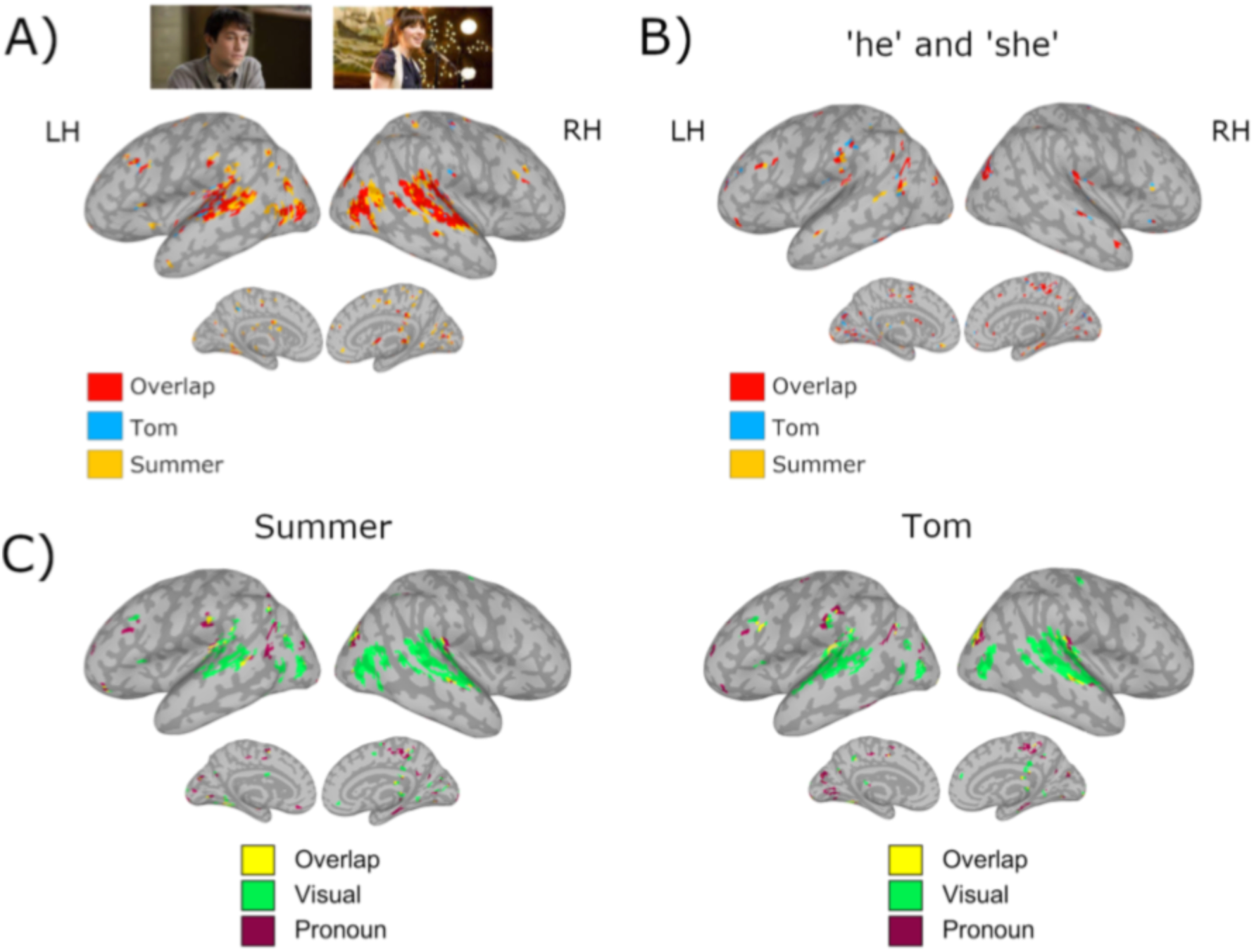
Saliency maps from vanilla guided backpropagation. For all figures, threshold = 95th percentile. A) Overlap of average visual maps for Tom vs Summer, where Red = overlap of two characters; Blue = Tom only; Yellow = Summer only. Cluster size =20. The two characters had high overlap in sensorimotor regions during visual references, with small differences in activity around these regions. B) Overlap of average pronoun maps for Tom vs Summer, where Red = overlap of two characters; Blue = Tom only; Yellow = Summer only. Cluster size =20. C) Overlap of average Summer and Tom maps for visual and pronoun references, where Yellow = overlap of two references; Green = visual only; Purple = pronoun only. Cluster size = 40. Pronoun and visual references overlapped in sensorimotor regions, with slightly different patterns

## Discussion

In this study, we tested the hypothesis that pronoun resolution reactivates unique sensorimotor character representations through context-dependent situation models, reconstructed during perceptual (i.e., visual) processing of a character in the same sensorimotor regions used during encoding.

To test this hypothesis, we implemented a 3D branched convolutional neural network that takes as input the 3D brain volumes of matched visual and pronoun references of a character in a movie. Our model achieved ∼93% accuracy in distinguishing the two main characters in a movie using both visual and pronoun references. Saliency maps thresholded at the 95^th^ percentile for the visual branch of the 3D bi-CNN model revealed that the model learned to distinguish between perceptual references of Tom and Summer through the bilateral involvement of primary and secondary visual regions, superior temporal gyri and sulci, parahippocampal and fusiform face area regions, and portions of primary motor cortex (Figure 3A). This distribution reflects our hypothesis that situation models are constructed via sensorimotor simulation.

Saliency maps thresholded at the 95^th^ percentile for the pronoun branch of the 3D bi-CNN model showed that the model used voxels in primary visual cortices, fusiform face area, hippocampus and parahippocampal area, precuneus, and mPFC to distinguish between Tom and Summer during pronoun resolution (Figure 3B). Our findings suggest that pronoun resolution requires access to the appropriate situation model to reactivate representations of a character, which was built and retrieved through sensory and motor activity.

### Model of pronoun resolution

Prior studies have proposed that narrative-based situation models may be built through sensory and motor activity (Zwaan, 2016), that are activated in the mPFC (Yarkoni et al., 2008), and left perisylvian regions during pronoun resolution (Hammer et al., 2011; Li et al., 2018). Further, it has been shown that schematic prior knowledge can help guide memory search during retrieval, with mPFC playing a particular role during encoding of these memories (Bero et al., 2014; Masis-Obando et al., 2022). However, aside from general aspects of encoding and retrieval processes, no concrete neuroimaging evidence thus far supports the existence of such simulations in the brain or suggests how pronouns may activate them.

Our findings show for the first time that understanding a reference to a particular individual occurs via active simulation involving situation models in brain regions involved in sensorimotor processing and theory of mind (“mentalising”). Based on our findings and the models already proposed (Altmann & Kamide, 2009; Wittenberg et al., 2021), we propose the following model of pronoun resolution based on previous visual information:

1. When a pronoun is uttered, brain networks focused on low level lexical processing engage with those focused on higher level semantic processing, specifically involving episodic memory retrieval, to activate a search in memory for the referent;
2. This search involves activating the appropriate situation model in which that referent is represented;
3. Once the appropriate situation model is active, the specific character (i.e., referent) is identified;
4. This character representation reactivates character-specific activity in sensorimotor regions, where the visual-based representation was originally constructed. Here we dissect some of these proposed processes.

### Pronouns and episodic memory

Imaging studies of pronoun resolution have identified a set of brain regions that activate when either (i) a referent is ambiguous (i.e., can refer to one of multiple characters); or (ii) the pronoun is incoherent with the antecedent (e.g., ”Julie was walking home. He had been at a party.“) (Hammer et al., 2007; Qiu et al., 2012). Such studies have found that pronoun resolution tasks activate primarily the IFG, MTG, and dorsolateral prefrontal cortex (dlPFC), with IFG activity associated with increasing task demands (e.g., ambiguity), MTG with incongruent gender of the pronoun and referent, and dlPFC having a general role in decision-making processes to help assign a referent (Hammer et al., 2007; Hammer et al., 2011; Hertrich et al., 2021; McMillan et al., 2012; Qiu et al., 2012).

In the work presented here, we did not find activation of MTG or IFG, with minimal activation of dlPFC (Figure 3B). Instead, pronouns for both the male and female character activated predominantly regions in the visual cortex (e.g., face fusiform area) and sensorimotor regions. The lack of activation in regions commonly invoked during language processing and typically engaged during incongruences between pronoun and referent may be due to the fact that the referents here were easily resolved through the rich contextual information afforded in the movie and commonly found in ecological discourse.

In this work, we found activation of a network composed of mPFC, hippocampus, parahippocampal area, and precuneus, likely involved in episodic memory and consolidation of context-dependent representations. Previous studies have suggested a fundamental role of mPFC in supporting situation model use (Baldassano et al., 2018; Yarkoni et al., 2008), likely through a role in networks commonly associated with mentalising and theory of mind, as well as the regions that are thought to be co-active during the “default mode". Here, the mPFC has a role in decision-making tasks, distinguishing between self and others and creating mental representations (Baetens et al., 2014; Cheetham et al., 2014; Isoda et al., 2013; Moran et al., 2011; Smith et al., 2018; Xu et al., 2005). However, as the mPFC was active along with the hippocampus, parahippocampal area and precuneus, its role is more likely to be in support of memory processes.

Previous research has shown that the mPFC is active during memory consolidation, particularly in long-term memory formation, after receiving information from the hippocampus (Euston et al., 2012; Takashima et al., 2006). The hippocampus is involved in short-term memory reactivation, notably during context-dependent episodic memory, together with the parahippocampal area and precuneus (Chang et al., 2021; Dickerson & Eichenbaum, 2010; Flegal et al., 2014; Maviel et al., 2004; Michelmann et al., 2021). Once information is fully consolidated, the mPFC inhibits activation of the hippocampus, to avoid building new representations of existing memories (Baldassano et al., 2018; Takashima et al., 2006). We thus propose that the activation of this network in the pronoun branch of our model includes two concurrent processes: (i) the hippocampus, parahippocampal area, and precuneus reinstate a situation model from working memory to help resolve the referent; and (ii) the mPFC updates the situation model with the newly encountered dialogue information and consolidates it in long-term memory. Given that the participants had not previously seen the movie, it is reasonable that the hippocampus would be engaged in the reactivation of context-specific situation models, with the mPFC consolidating these representations. Possible extensions of this work could examine possible temporal variations in hippocampus/ mPFC over the movie, as well as testing this activity in repeated movie viewings.

### Situation models and character representations

Although much of the literature on situation models has focused on building discourse representations (Zwaan, 2016; Zwaan et al., 1995; Zwaan & Radvansky, 1998), to the best of our knowledge, there is still no biological evidence for their representation in the brain or for their involvement in pronoun resolution. Here we demonstrated the existence of context-dependent situation models for the representation of characters and their reactivation during pronoun resolution. In the findings discussed here, situation models containing character representations are generally constructed in regions of the brain mediating sensory and motor processing (Figure 3A).

Brain activation for resolving the two main characters of “500 Days of Summer” primarily involved voxel distributions in and around the primary visual cortices, occipitotemporal and occipitoparietal regions, and superior temporal gyri and sulci in both hemispheres (Figure 3A). These regions are involved in visual perception via both dorsal and ventral streams (Ungerleider & Mishkin, 1982): The ventral visual pathway initially forms different distributions in the primary visual cortex that relate to face perception of different characters (Sheth & Young, 2016), and then the dorsal pathway engages the occipitoparietal area to process information visually encoded by actions (Freud et al., 2016). The superior temporal gyrus and sulcus have recently been shown to form a third pathway that integrates the previous two to process social interactions afforded by visual information (e.g., gestures, facial expressions) (Manfredi et al., 2017; Pitcher & Ungerleider, 2021).

The character representations in the brain are not solely due to visual perception, although regions important in visual perception are certainly integral: Additional activation included regions known to play a role in imagery, abstraction, attention, and construction of representations, as illustrated by the Neurosynth meta-analysis term correlations (see Table 1). Given that situation models are highly imagistic by nature (Zwaan, 2016), the presence of clusters relating to imagery further suggest that the 3D bi-CNN model has isolated processes/regions involved in building situation models of character representations.

**Table 1.** Top 5 Neurosynth functional terms which correlated to six clusters in each character’s visual and pronoun reference. The visual references mapped mainly to sensorimotor regions involved in visual, motor and language processing, but also included some elements of imagery/abstraction. These patterns of activity were reactivated during pronoun references. The latter also activated regions associated with episodic memory and mentalising.

|  |  | Cluster 1 | Cluster 2 | Cluster 3 | Cluster 4 | Cluster 5 | Cluster 6 |
| --- | --- | --- | --- | --- | --- | --- | --- |
| **Summer Visual** | **Term 1** | Motion | Sentences | Motion | Execution | Visual | Imagery |
|  | **Term 2** | Video | Languages | Visual | Movement | Eye Movements | Objects |
|  | **Term 3** | Action Observation | Construction | Action Observation | Motor Imagery | Video | Finger Tapping |
|  | **Term 4** | Perception | Word | Videos | Executive Functions | Attention | Navigation |
|  | **Term 5** | Visual | Syntactic | Gestures | Task | Imagery | Abstract |
| **Summer Pronoun** | **Term 1** | Mentalizing | Task | Eye | Visual | Place | Episodic |
|  | **Term 2** | Social | Demands | Visual | Likelihood | Objects | Autobiographical Memory |
|  | **Term 3** | Theory Mind | Working | Colour | Reading | Navigation | Retrieval |
|  | **Term 4** | Default | Language | Visuo Spatial | Language Network | Face | Experiences |
|  | **Term 5** | Person | Phonological | Retrieval | Abstract | Encoding | Construction |
| **Tom Visual** | **Term 1** | Motion | Sounds | Objects | Hearing | Visual | Movements |
|  | **Term 2** | Visual | Speech | Face | Listening | Location | Finger Tapping |
|  | **Term 3** | Action Observation | Listening | Item | Vocal | Object | Hand |
|  | **Term 4** | Videos | Pitch | Visual | Sensory | Concrete | Coordination |
|  | **Term 5** | Perception | Voice | Decision Task | Noise | Abstract | Motor Task |
| **Tom Pronoun** | **Term 1** | Referential | Theory of Mind | Audiovisual | Speech | Place | Efficiency |
|  | **Term 2** | Thoughts | Navigation | Preparatory | Speaker | Objects | Multisensory |
|  | **Term 3** | Default Network | Visual | Abstract | Painful | Navigation | Abstract |
|  | **Term 4** | Theory of Mind | Virtual | Abuse | Sound | Face | Abuse |
|  | **Term 5** | Personality Traits | Abstract | Accurate | Acoustic | Encoding | Accurate |

### Content retrieval

Our findings support the hypothesis that pronouns reactivate sensorimotor “fingerprints” related to the character representations in different situation models. We found that the primary visual cortex and parts of the occipitotemporal area, occipitoparietal cortex, and superior temporal gyrus and sulcus were reactivated during pronoun resolution (Figure 3B, C). These suggest that when a pronoun is uttered, a search for the situation model in working memory reinstates the activity distribution specific to the representation of the referent.

Studies on content retrieval have shown that the brain reactivates specific perceptual activity fingerprints when recalling distinct contextual information in the absence of the stimulus. Indeed, there is ample evidence from the study of episodic memory that retrieval success correlates with the degree of reactivation of activity patterns from antecedent to new event (Frankland et al., 2019; Oedekoven et al., 2017; Yaffe et al., 2014). For instance, a study using naturalistic stimuli in the form of short videos showed that increased overlap between antecedent and reinstatement increases vividness of the antecedent video during retrieval (St-Laurent et al., 2015). The patterns reactivated during free recall are unique to the category they represent (e.g., face, object, place) (Polyn et al., 2005) or the context to which they refer (Nyberg et al., 2000), although overlap between activity distributions of categories/ contexts proportionally increases with their similarity (Norman et al., 2006). In line with previous research, we found that the sensorimotor activity distributions of representation of characters Tom and Summer mostly overlapped, unsurprising given their shared membership in the ‘face’ category or similar contextual situation models (Figure 3). Future work might explore the differences in these patterns using a representation similarity analysis or analogous method.

Our study builds upon existing knowledge about content retrieval, by linking visual information of specific characters to their unique retrieval through pronouns in a naturalistic setting, where the stimuli are complex and continuous and free recall cannot be tested.

### Limitations

One limitation of the present study relates to the genders of the movie’s two main characters, i.e., one male and the other female. This certainly made the anaphora resolution task a bit easier.

A second limitation relates to the limited number of pronouns in movies, and our decision not to remove those pronouns occurring during a flashback or during an on-screen presence of a character referent. For example, Summer’s face appears in some scenes where other characters are referring to her as ‘she’, and similarly, Tom may appear when the narrator in the movie refers to him as ‘he’. This limitation may bias the model towards a character’s face rather than the pronoun retrieval model. Nevertheless, this occurred in only 29% of the 84 initially detected pronouns (n.b., some pronouns referring to Tom and Summer may not be detected by the speech-to-text transcript and thus would not be included as samples here). Moreover, these instances usually occurred as single blocks, and by excluding pronouns less than 5 seconds apart in either past or future directions (resulting in 21 pronoun samples for Summer and 8 for Tom prior to balancing), we further reduced the co-occurrence of characters on-screen. To improve on this potential confound in future work, we could ignore instances where the face is on screen during pronoun reference and make use of additional movies and participants to obtain a larger number of samples for training/testing.

Finally, a third limitation of this study lies in the selection of the machine learning architecture, specifically the use of a shallow CNN comprising a single convolutional and fully connected layer. While computationally efficient, such a model may lack the representational depth necessary to capture the complex neural patterns associated with pronoun processing. This constraint, combined with the limited size of the training and test datasets, could have led to overfitting that may not be detectable due to the small sample size, despite the described steps being taken to minimise this possibility. Given that pronoun-related activations are distributed across multiple brain regions, more expressive models would be needed to extract meaningful features. In future work, we intend to fine-tune a pre-trained 2D ResNet model adapted for 3D MRI data, which could offer improved generalizability, more robust signal detection, and superior capture of the spatial complexity inherent in the task.

### Implications

The present study demonstrates that pronoun resolution requires reactivation of unique character representations from perceptually built situation models. In particular, we showed that (i) situation models for character representations are encoded in the brain, and (ii) performing anaphoric reference invokes sensorimotor character representations.

This study has significant implications for understanding the neurobiology of language in the real world. Existing models generally do not account for linguistic complexities and how these may drive multi-modal interactions between language, memory, and perception, which are known to be elicited by context (Friston & Price, 2001; Skipper, 2015). This study demonstrated that when inspecting natural contextual dependencies in specific linguistic features, the distribution of activity includes regions outside classical ‘language’ areas and requires the interplay between various modalities (Aliko et al., 2023).

Difficulties in pronoun resolution are a common feature of aphasia, irrespective of language (Arslan et al., 2021), and particularly in agrammatic aphasia (Jarema & Friederici, 1994). Importantly, some people with aphasia have difficulty associating the pronoun to the correct subject referent, highlighting how the process of retrieval may be impaired (Peristeri & Tsimpli, 2013). Individuals with Alzheimer Disease (Almor et al., 1999) and Mild Cognitive Impairment (Lust et al., 2024) have significant problems with anaphora resolution. Healthy older adults also show a decrease in this ability, particularly with increasing amounts of material intervening between the pronoun and the referent (Light & Capps, 1986). Our finding that pronoun resolution depends on the reactivation of sensorimotor character representations could offer insights for the development of novel therapies that target more specific language features and associated processes to improve performance in these situations.

## Conclusion

In this study we demonstrated that pronouns in naturalistic discourse reactivate a set of sensorimotor character representations incorporated into situation models constructed when the characters were visually present. These schematic memory representations are built using both perceptual and more general cognitive processes, suggested by recruitment of regions important for “mentalising” and “default mode” processing required for forming imagistic simulations. The sensorimotor distribution of character representations mostly overlapped between the two characters, suggesting that they shared similar context in the movie. However, character representations manifested small variations around visual regions, demonstrating that situation models can help point to a unique character representation during pronoun resolution.

Overall, these findings highlight the importance of studying individual language features in a more complex and natural environment and demonstrate that the neurobiology of language is more distributed than existing models suggest (Aliko et al., 2023; Drijvers et al., 2025). The distributed nature of these representations may offer new avenues for treatment of individuals with aphasia.

## Acknowledgements

This work was partially supported by a doctoral training grant given to SA from the UK Research and Innovation (UKRI) Biotechnology and Biological Sciences Research Council (BBSRC) as part of the London Interdisciplinary Biosciences Consortium (LIDo). We would like to thank the Birkbeck-UCL Centre for Neuroimaging (BUCNI) for supporting the NNDb. We thank Bangjie Wang for technical assistance, Annette Glotfelty for helpful comments on the manuscript, and the UCL LAB Lab and the UTD Brain Circuits Lab for informative discussion.

## Conflicting Interests

The authors declare that they have no conflicts of interest.

## Open Science Statement

This study was not preregistered. A version of this work is available in SA’s Doctoral thesis, ‘A Model of the Network Architecture of the Brain that Supports Natural Language Processing’. The preprocessed data used in analyses is publicly available on OpenNeuro (https://openneuro.org/datasets/ds002837/versions/2.0.0). The code used to generate results will be made available on GitHub (https://github.com/lab-lab/).

## Notes

https://openneuro.org/datasets/ds002837/versions/2.0.0

### Competing Interest Statement

The authors have declared no competing interest.

