## Supplementary Figures for "Situation Models in the Brain are Used to Resolve Word References"

#### *Figure S1*

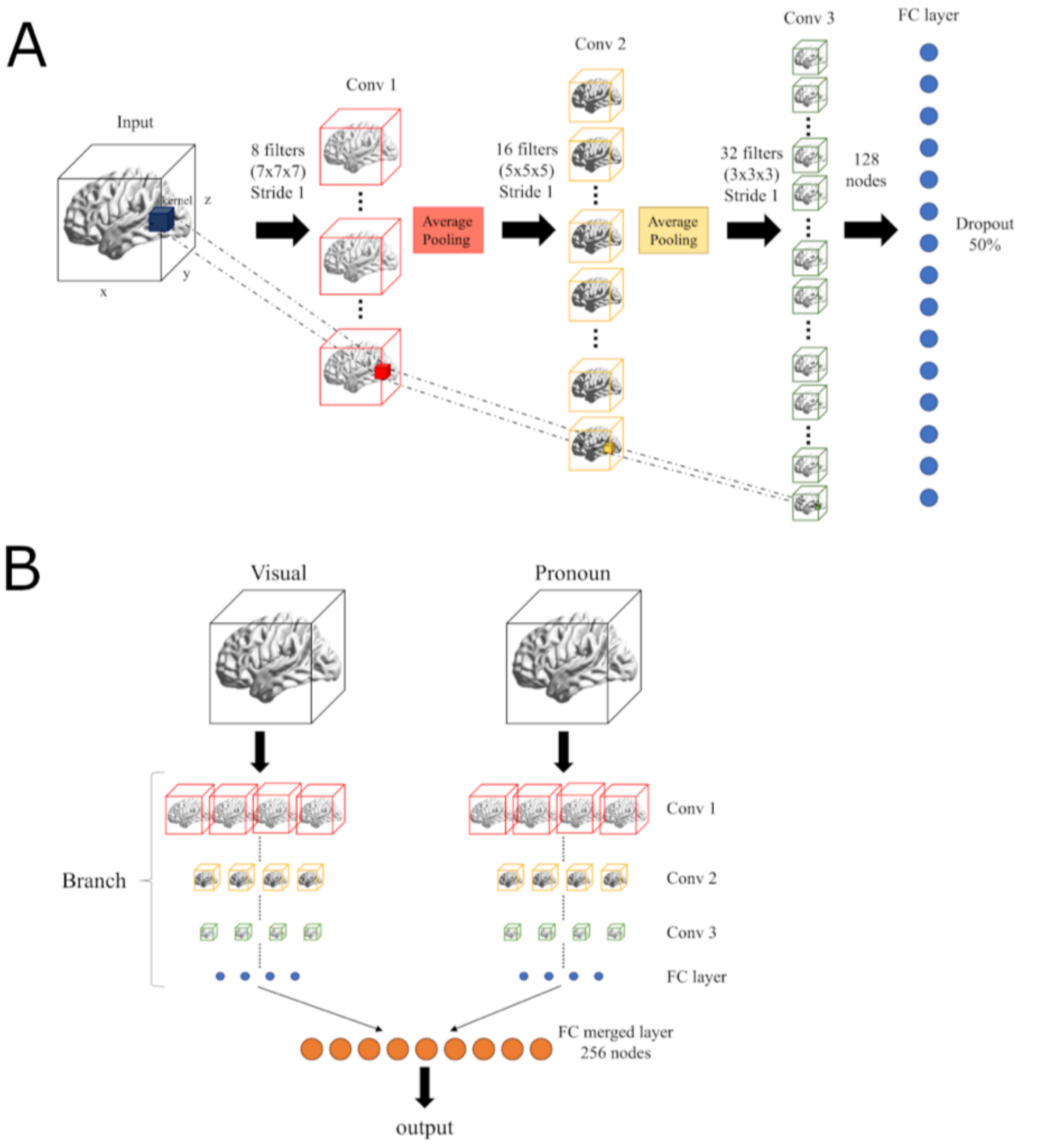


**Figure S1**. (A) Schematic of the 3D CNN from Vu et al. (2020). A single MRI volume is processed with 7×7×7 kernels (8 filters) and pooling (layer 1), followed by 5×5×5 kernels (16 filters) with pooling (layer 2), and 3×3×3 kernels (32 filters, layer 3). The output is passed to a fully connected layer (128 nodes) for classification.

(B) Schematic of the 3D bi-CNN architecture with visual and pronoun branches, each using the structure in (A). Outputs from the branches are concatenated and passed to a fully connected layer (256 nodes) before classification.

*Figure S2*

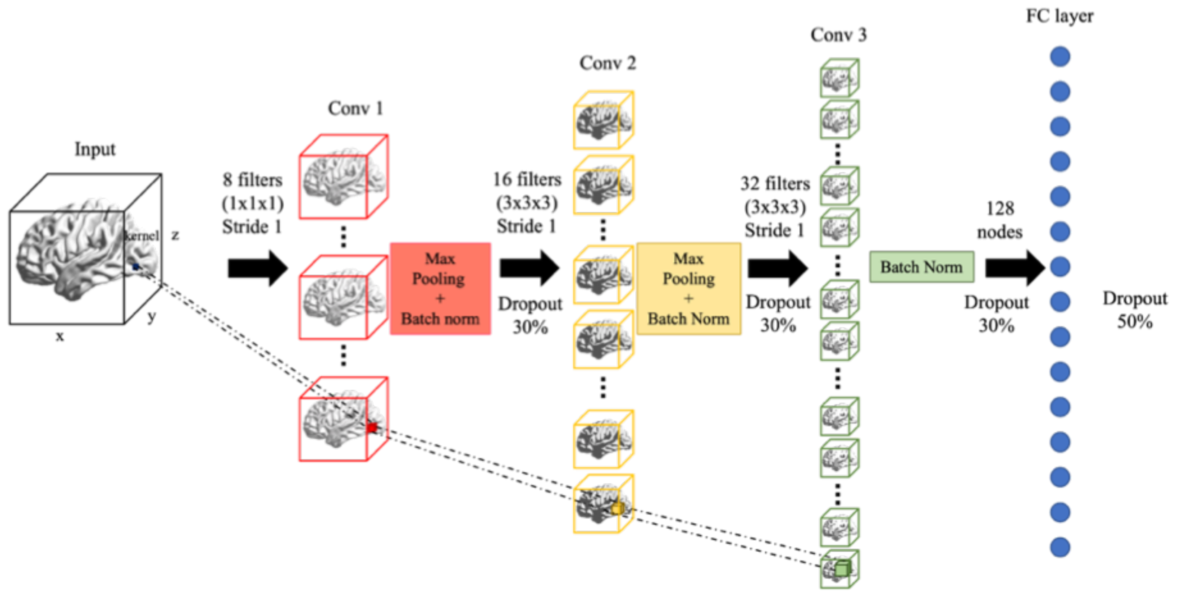


**Figure S2**. Modifications to Vu et al (2020) architecture: (1) layer 1 kernel reduced to 1×1×1, (2) average pooling replaced with max pooling and batch normalisation, (3) layer 2 kernel reduced to 3×3×3, (4) batch normalisation added before the fully connected layer, (5) 30% dropout applied after each convolutional layer.

### *Figure S3*

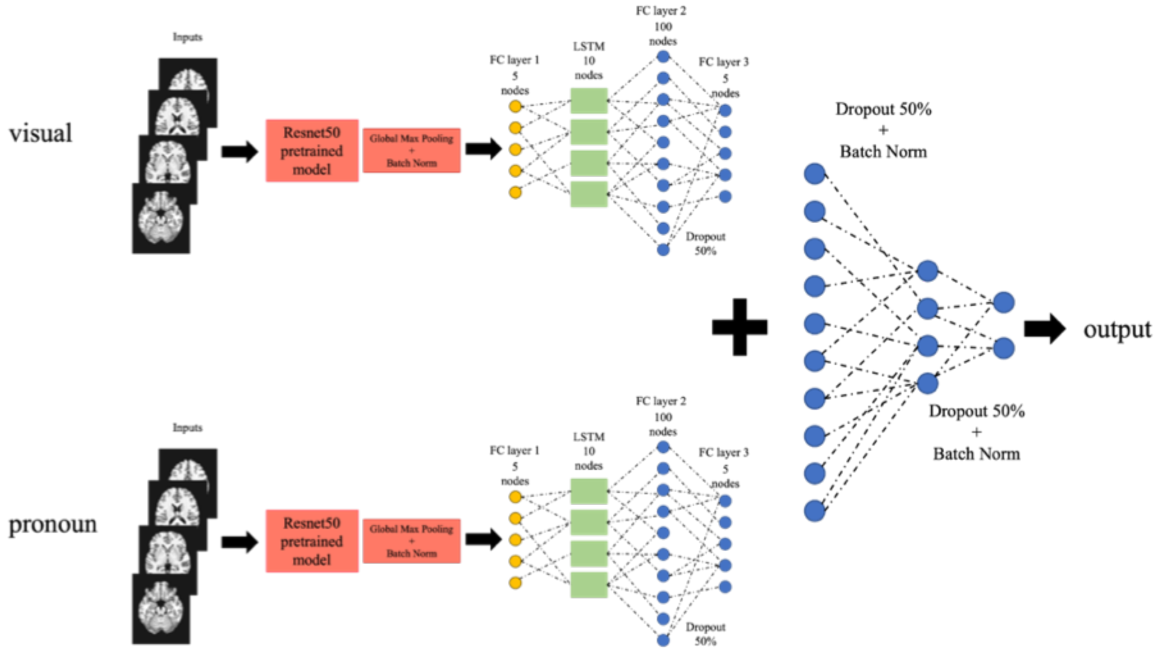


**Figure S3**. Schematic of the 2D CNN branched architecture. Axial MRI slices are input to pretrained ResNet50 models for pronoun and visual branches. Outputs are pooled, batch normalised, and passed to a 5-node fully connected layer. An LSTM (10 nodes) maintains slice order, followed by two fully connected layers per branch. Branch outputs are merged and passed through two fully connected layers with 50% dropout and batch normalisation before classification.
